# Genetic surveillance of SARS-CoV-2 M^pro^ reveals high sequence and structural conservation prior to the introduction of protease inhibitor Paxlovid

**DOI:** 10.1101/2022.03.29.486331

**Authors:** Jonathan T. Lee, Qingyi Yang, Alexey Gribenko, B. Scott Perrin, Yuao Zhu, Rhonda Cardin, Paul A. Liberator, Annaliesa S. Anderson, Li Hao

## Abstract

SARS-CoV-2 continues to represent a global health emergency as a highly transmissible, airborne virus. An important coronaviral drug target for treatment of COVID-19 is the conserved main protease (M^pro^). Nirmatrelvir is a potent M^pro^ inhibitor and the antiviral component of Paxlovid™. The significant viral sequencing effort during the ongoing COVID-19 pandemic represented a unique opportunity to assess potential nirmatrelvir escape mutations from emerging variants of SARS-CoV-2. To establish the baseline mutational landscape of M^pro^ prior to the introduction of M^pro^ inhibitors, M^pro^ sequences and its cleavage junction regions were retrieved from ∼4,892,000 high-quality SARS-CoV-2 genomes in GISAID. Any mutations identified from comparison to the reference sequence (Wuhan-hu-1) were cataloged and analyzed. Mutations at sites key to nirmatrelvir binding and protease functionality (e.g., dimerization sites) were still rare. Structural comparison of M^pro^ also showed conservation of key nirmatrelvir contact residues across the extended *Coronaviridae* family (alpha-, beta-, and gamma-coronaviruses). Additionally, we showed that over time the SARS-CoV-2 M^pro^ enzyme remained under purifying selection and was highly conserved relative to the spike protein. Now, with the EUA approval of Paxlovid and its expected widespread use across the globe, it is essential to continue large-scale genomic surveillance of SARS-CoV-2 M^pro^ evolution. This study establishes a robust analysis framework for monitoring emergent mutations in millions of virus isolates, with the goal of identifying potential resistance to present and future SARS-CoV-2 antivirals.

**Importance:** The recent authorization of oral SARS-CoV-2 antivirals, such as Paxlovid, has ushered in a new era of the COVID-19 pandemic. Emergence of new variants, as well as selective pressure imposed by antiviral drugs themselves, raise concern for potential escape mutations in key drug binding motifs. To determine the potential emergence of antiviral resistance in globally circulating isolates and its implications for the clinical response to the COVID-19 pandemic, sequencing of SARS-CoV-2 viral isolates before, during, and after the introduction of new antiviral treatments is critical. The infrastructure built herein for active genetic surveillance of M^pro^ evolution and emergent mutations will play an important role in assessing potential antiviral resistance as the pandemic progresses and M^pro^ inhibitors are introduced. We anticipate our framework to be the starting point in a larger effort for global monitoring of the SARS-CoV-2 M^pro^ mutational landscape.

## Introduction

The causative agent of coronavirus disease 2019 (COVID-19) was identified as a novel coronavirus (CoV) (1), later named severe acute respiratory syndrome coronavirus 2 (SARS-CoV-2), with close genetic and clinical resemblance to the 2002 SARS virus (SARS-CoV) (2, 3). SARS-CoV-2 shares the core features of all CoVs, including a large positive-stranded RNA genome (26-32 kb), the spike (S), envelope (E), membrane (M), and nucleocapsid (N) structural proteins, as well as two conserved viral proteases: the main protease (M^pro^), also known as 3-chymotrypsin-like cysteine protease (3CL^pro^), and papain-like protease (PL^pro^) (4). These enzymes digest two large polyproteins (pp1a and pp1ab) at multiple junctions to generate a series of proteins critical for virus replication and transcription, including the RNA-dependent RNA polymerase (RdRp), helicase, and the M^pro^ protein itself (5). M^pro^ is encoded by ORF1 as non-structural protein 5 (Nsp5) and cleaves the polyproteins at 11 sites to release Nsp4-Nsp16, making M^pro^ an essential protein for the CoV life cycle (6).

Since the onset of the COVID-19 pandemic in 2020, SARS-CoV-2 variants have rapidly emerged worldwide, raising concern for the effectiveness of currently available vaccines and neutralizing monoclonal antibodies (mAbs) targeting the S protein. As of March 2022, the World Health Organization (WHO) has identified five major variants of concern (VOCs): B.1.1.7 (Alpha, α), B.1.351 (Beta, β), P.1 (Gamma, γ), B.1.617.2 (Delta, Δ) and most recently, B.1.1.529 (Omicron, ο) (7). Characterization of emergent variants has centered on the number and location of mutations in the S protein trimer (8). Omicron, specifically, contains several signature mutations in the S protein that enable the variant to escape immunity from previous infection or vaccination (9), making it unlikely that each of the approved mAbs will maintain clinical efficacy against this VOC (10). To date, the only approved or authorized non-mAb therapeutics for COVID-19 are small-molecule antivirals: remdesivir and molnupiravir, both RdRp inhibitors originally developed for different RNA viruses, and Paxlovid™, whose antiviral component, nirmatrelvir, a CoV M^pro^ inhibitor, is co-administered with ritonavir. Remdesivir is administered intravenously, while molnupiravir and Paxlovid are orally bioavailable.

Nirmatrelvir is an active site inhibitor of the SARS-CoV-2 M^pro^ that exhibits *in vitro* antiviral activity across the *Coronaviridae* family, demonstrating potent inhibition of the M^pro^ from all other beta-coronaviruses (β-CoVs) and alpha-coronaviruses (α-CoVs) known to infect humans (11). Active sites of M^pro^ are largely conserved among β-CoVs. The SARS-CoV-2 M^pro^ amino acid sequence shares 96% identity to that of SARS-CoV, with differences at 12 residues between the two viruses (12). The critical amino acid residues involved in enzyme-inhibitor binding interactions are also particularly well-conserved within this family of viruses (13). Its essential functional importance in virus replication, together with the absence of closely related homologues in humans (14), identify the CoV M^pro^ as an attractive antiviral drug target (11, 15). Indeed, Paxlovid was granted Emergency Use Authorization (EUA) from the US Food and Drug Administration (FDA) in December 2021, after positive results in the Phase 2/3 EPIC-HR (Evaluation of Protease Inhibition for COVID-19 in High-Risk Patients) trial (16).

In such a rapidly evolving pandemic, it is important to monitor resistance of emerging variants to compounds targeting critical viral proteins, including M^pro^. Among the many unprecedented aspects of the ongoing COVID-19 pandemic is an intense phylogenetic surveillance of the virus in the human population. The genome sequence of millions of SARS-CoV-2 isolates has been determined and deposited into the open-access Global Initiative on Sharing Avian Influenza Data (GISAID) database (17) since January 10, 2020. The accessibility of real-world sequences from the expansive GISAID dataset has enabled a global, collaborative effort by scientists to track emerging lineages, identify signature escape mutations, and classify new variants in real time (18). To our knowledge, a comprehensive genomic surveillance of mutations in SARS-CoV-2 nonstructural proteins is limited to the RdRp (19, 20). Large-scale genetic surveillance of the M^pro^ enzyme from circulating SARS-CoV-2 variants has yet to be reported.

In the present study, we built a workflow to monitor the evolution of M^pro^ and the emergence of potential escape mutations in millions of SARS-CoV-2 genomes obtained from GISAID. We address the suitability of M^pro^ as a drug target for COVID-19 by evaluating polymorphisms at M^pro^ dimerization and substrate cleavage sites, in addition to key contact residues with the selective inhibitor, nirmatrelvir, and thus provide a baseline understanding of M^pro^ diversity prior to the widespread use of Paxlovid.

## Materials and Methods

### Structural comparison of M^pro^ from different CoVs

The crystal structures of M^pro^ from multiple CoVs have been reported previously in either apo or inhibitor-bound form (21–24). The PDB structures selected as representatives for analysis are listed in Table S1 (*n=*12). The active site amino acids are defined as those within 4.5Å of the common ligand PRD_002214. The chain A of 11 M^pro^ proteins were superimposed on the SARS-CoV-2 M^pro^ protein complexed with nirmatrelvir (PDB 7RFW) based on the Carbon-α (Cα) of the 26 amino acids. The superposition of images was generated using the Molecular Operating Environment (MOE) software platform (version 2020.09, Chemical Computing Group ULC, Montreal, QC, Canada). The root mean square deviation (RMSD) was also calculated based on the 26 Cα atoms.

### SARS-CoV-2 genomes and M^pro^ annotation pipeline

Genome sequences and patient metadata for ∼4.9 million isolates were obtained from the GISAID (17) EpiCoV database (www.epicov.org) through January 14, 2022. The genomes were quality filtered: incomplete genomes <29,000 nucleotides in length and/or containing >5% ambiguous nucleotides (Ns) were excluded. Sequences, collection dates, countries of origin, and lineage assignments were deposited to an internal database, BIGSdb (25).

M^pro^ nucleotide sequences were obtained using BLASTN alignment (26) to the reference SARS-CoV-2 genome (NC_045512.2, isolate Wuhan-hu-1) (27). Sequences with less than 90% alignment or containing ambiguous bases were excluded from further analysis. Nucleotide alleles were translated to amino acid sequences and non-synonymous polymorphisms were called through pairwise alignment to the reference M^pro^ amino acid sequence of the Wuhan-hu-1 isolate. Protein sequences were assigned unique ID linked to the respective viral genomes in BIGSdb.

### Non-synonymous mutation rate calculation

A list of mutation fingerprints (MF) was downloaded from COVID-19 Virus Mutation Tracker (CoVMT) (28) (https://www.cbrc.kaust.edu.sa/covmt/). MF was defined as the specific set of mutations shared by a group of genomic isolates from GISAID. This list is regularly updated and maintained by the CoVMT team. An ad-hoc script was written to calculate the number of non-synonymous mutations occurring on the M^pro^, RdRp and S genes per month. The amino acid mutation rate for each gene was then calculated and plotted by month of sample collection.

### Nucleotide diversity and *d*_N_/*d*_S_ selection analysis

Because selection analysis tools are computationally intensive, the genome dataset retrieved from GISAID was randomly downsampled to a manageable subset (∼25K) using the Nextstrain Augur pipeline (29) with a maximum of 30,000 sequences equally sampled by geographic region and month from December 1, 2020 through January 1, 2022. Three downsampled subsets of SARS-CoV-2 genomes were independently generated. Each subset of genomes was then aligned to the reference genome (Wuhan-hu-1) using MAFFT (30) (with –6mer pair flag for rapid alignment of large numbers of closely related viral genomes). M^pro^, RdRp and S genes were extracted from the genome-wide alignments. To prepare for selection analysis, sequences with entries of N or with deletions (-) were filtered out and STOP codons were replaced with triplet of (-). As the S gene has many deletions, to maintain a comparable number of sequences, the sequences with deletions were not filtered out, and instead, those with non-in-frame deletions were replaced with in-frame deletions. Overall nucleotide diversity was inferred using MEGA X (31). The ratio of non-synonymous-to-synonymous mutations (***d*_N_/*d*_S_** or ω) was inferred using GenomegaMap (32) (Bayesian sliding window model) with the transition:transversion ratio (κ) of 1.0 and nucleotide diversity (θ) of 0.17. Two independent Markov chain Monte Carlo (MCMC) analyses were run at 500,000 iterations each. Runs were compared for convergence and the resulting ***d*_N_/*d*_S_** determined using RStudio (version 1.1.383). The average of ***d*_N_/*d*_S_** from three downsampled datasets were used for our selection analysis.

### SARS-CoV-2 intra-lineage M^pro^ diversity analysis

The five most prevalent M^pro^ protein sequences among GISAID isolates were retrieved from BIGSdb for each variant of interest (VOI) or VOC. Any polymorphisms among these sequences were determined from the prior alignments. The total instances of each mutation were then obtained based on sequence prevalence within each SARS-CoV-2 lineage.

### Structural analysis of the M^pro^ dimer interface

Residues involved in stabilization of the M^pro^ dimer interface were identified from the structure of the dimeric SARS-CoV-2 M^pro^ 7RFR.pdb (11) (Table 1). Interprotomer contacts were initially identified using Biovia Discovery Studio Visualizer (version 4.5, Dassault Systèmes) and then manually inspected to confirm. All structural models of the M^pro^ protein were rendered using the Biovia Discovery Studio Visualizer software.

**Table 1.**
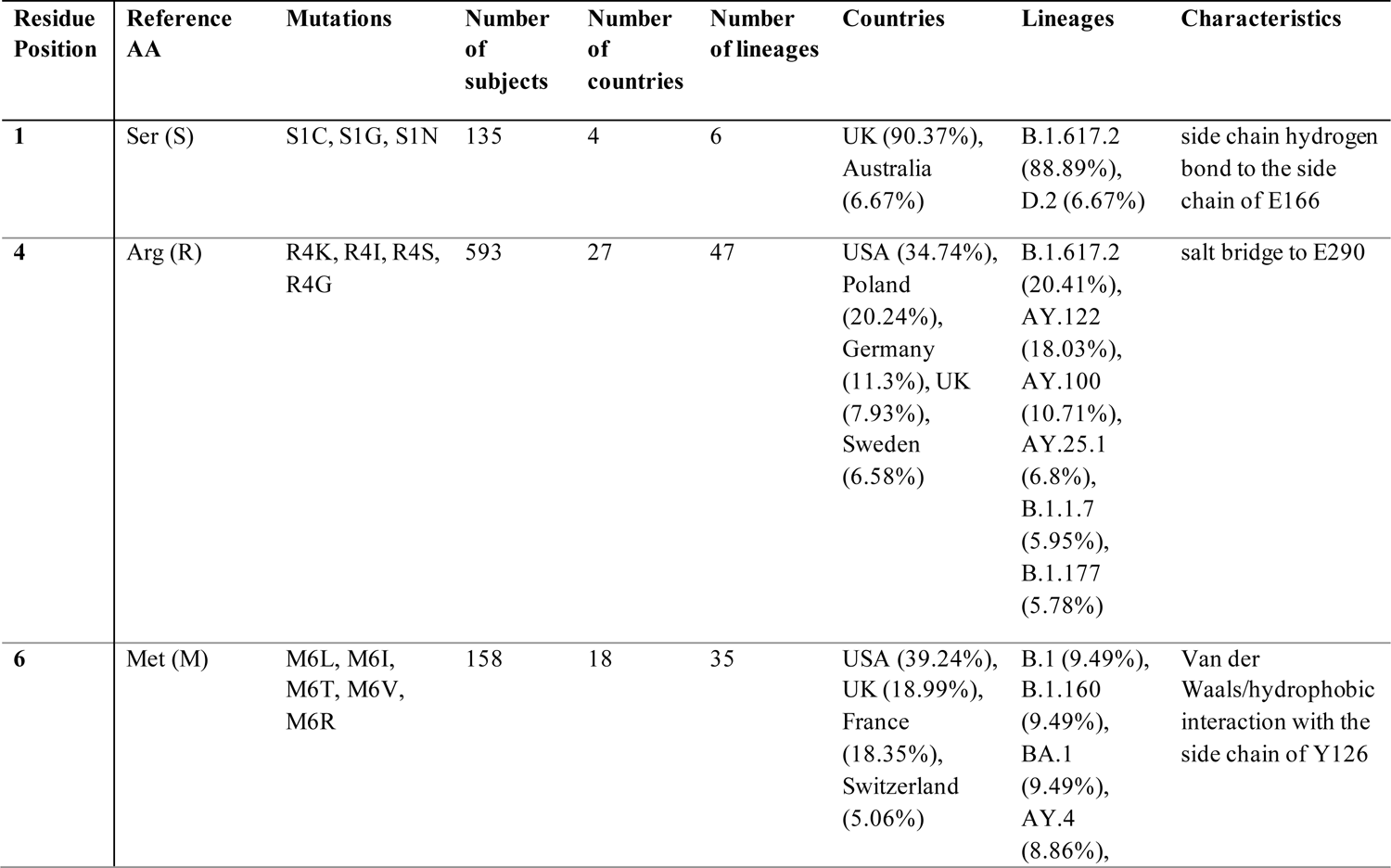
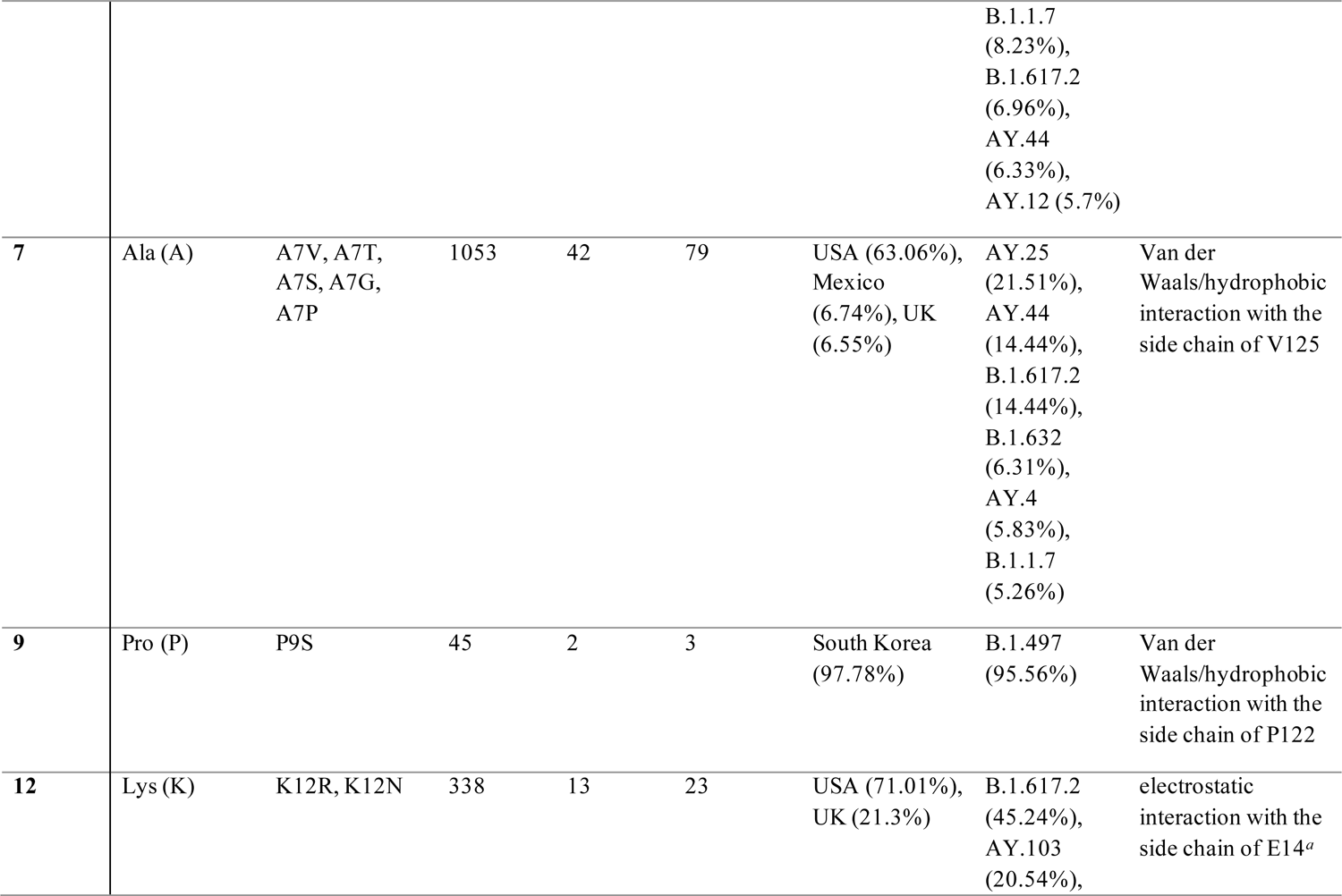
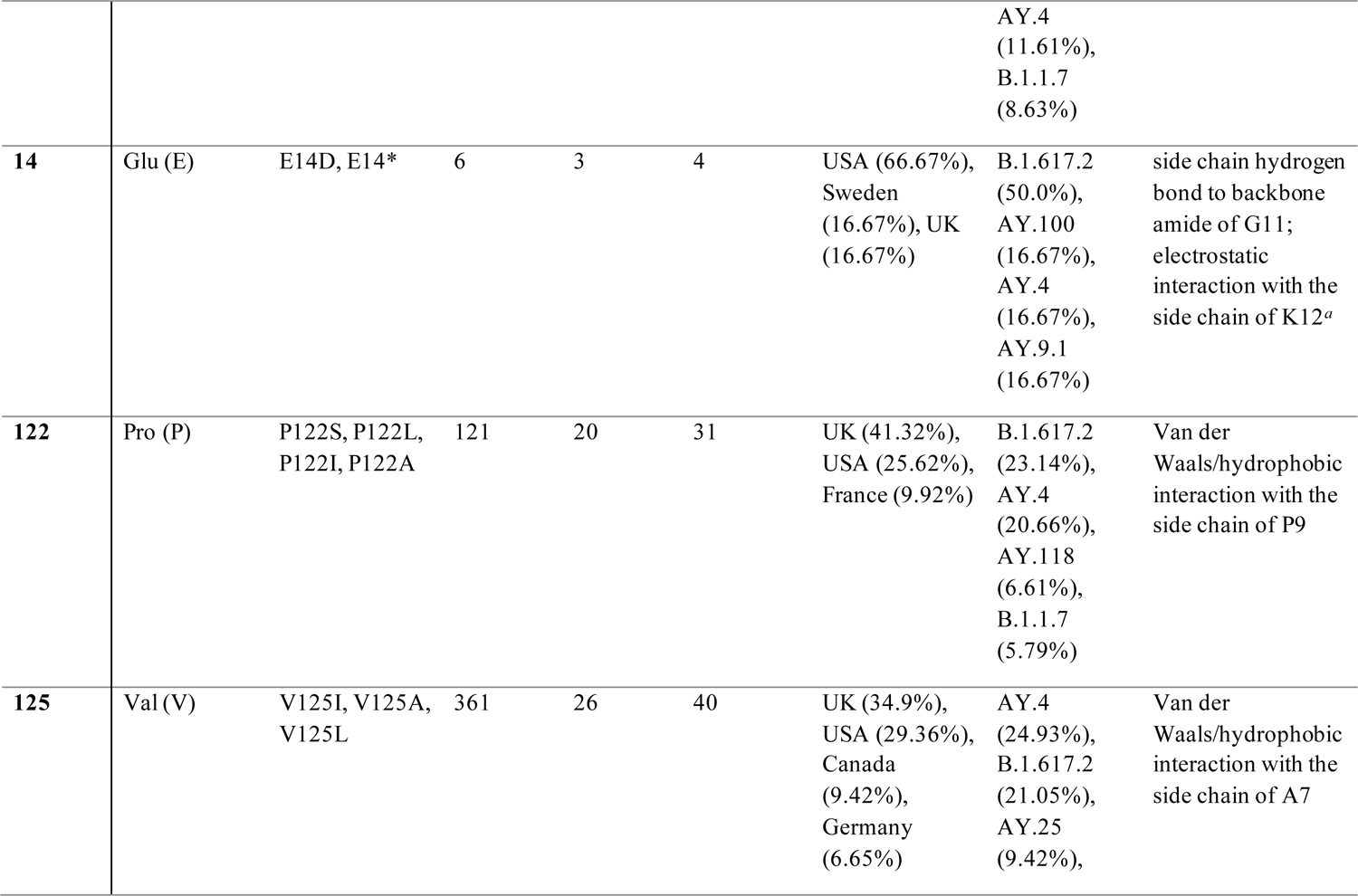
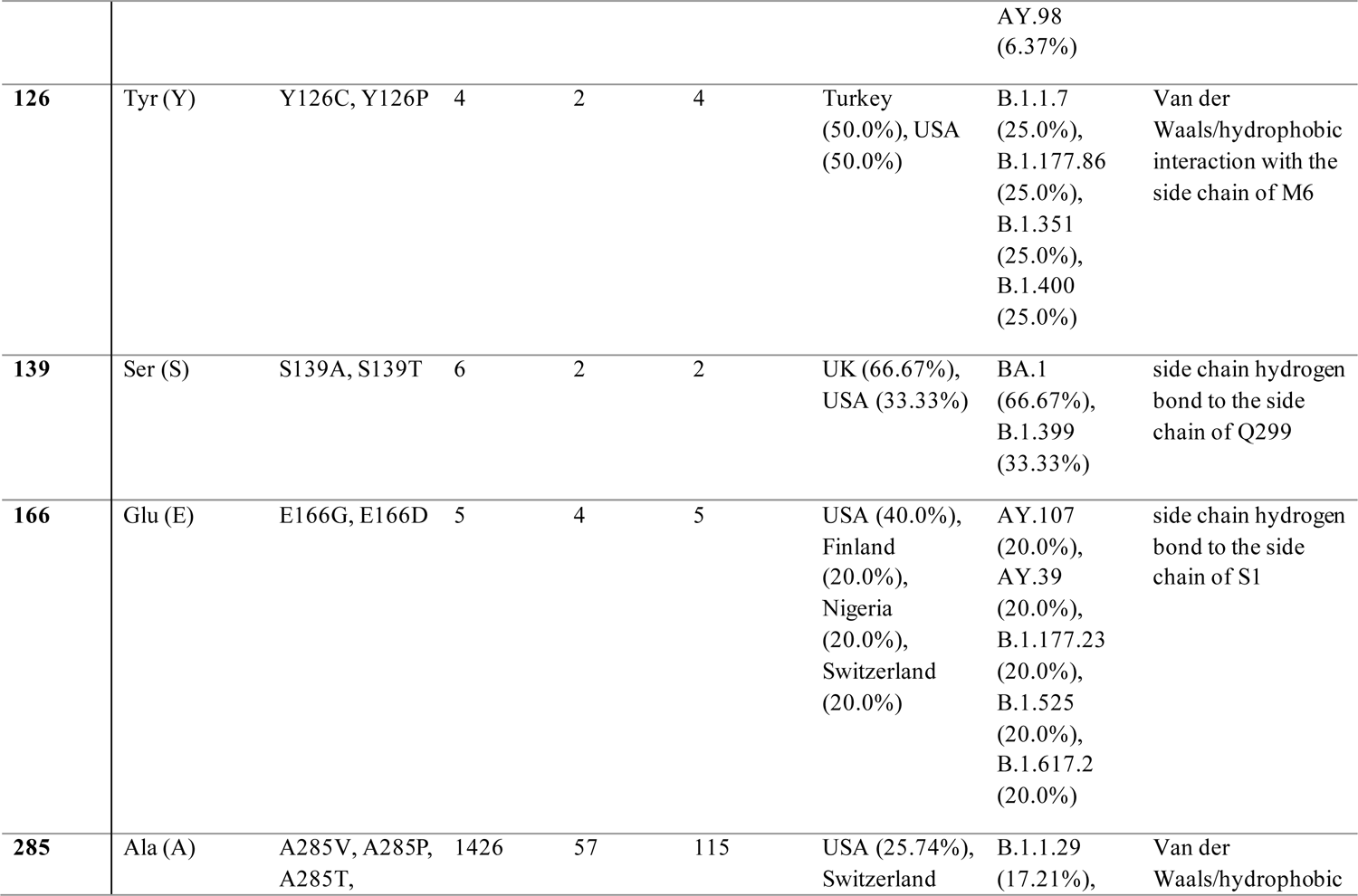
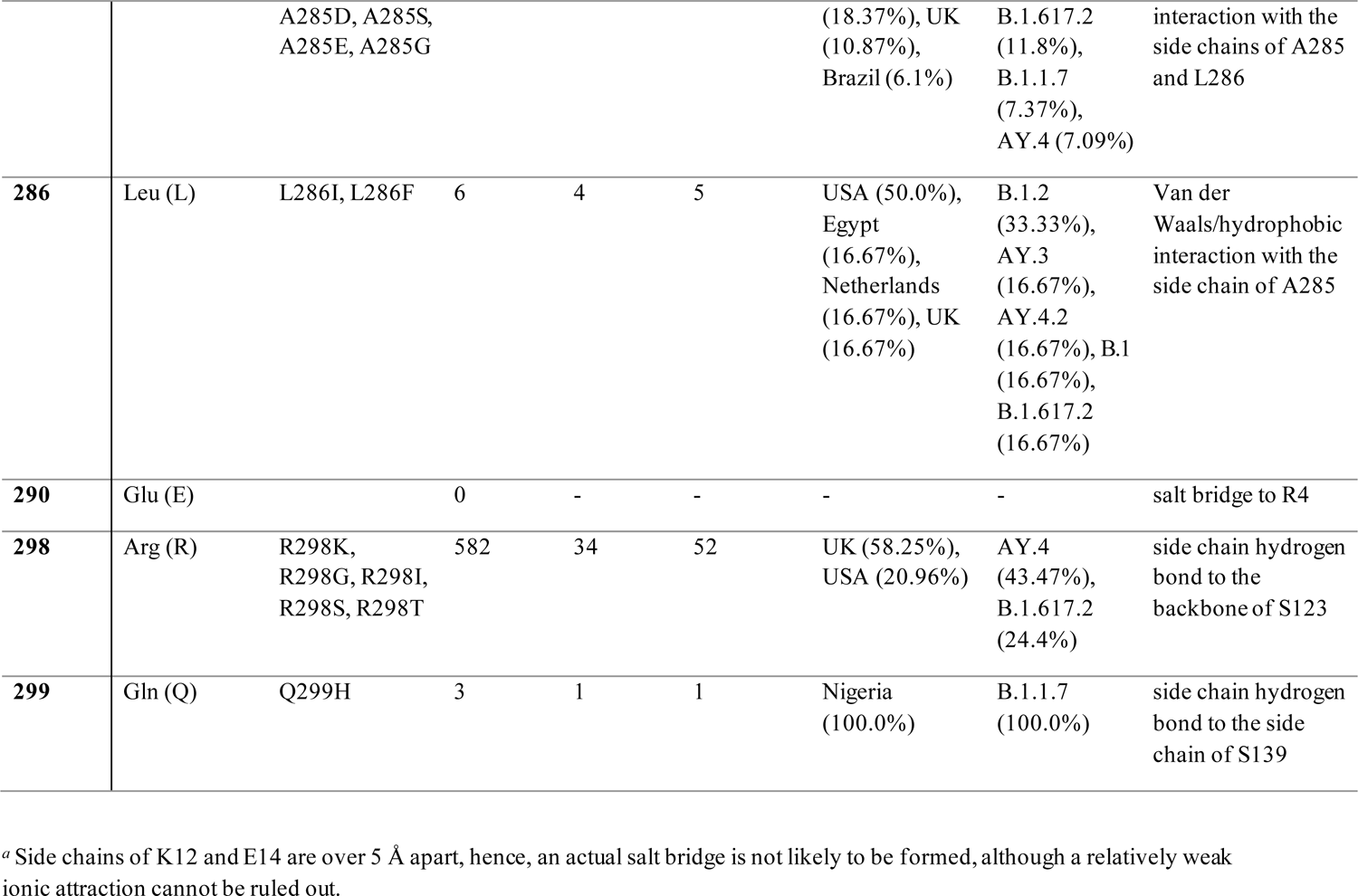
Mutation breakdown at M^pro^ dimerization interface residues.

### Data availability

All viral genome sequences analyzed herein were obtained from the GISAID public database (17) (www.gisaid.org). These sequences represented accessions for samples deposited between January 10, 2020 and January 14, 2022. The accession numbers total in the millions.

## Results

### Structural and sequence conservation of M^pro^ from different CoVs

Nirmatrelvir was previously demonstrated to have robust pan-CoV antiviral activity (11). To further investigate the conservation of M^pro^ across the extended *Coronaviridae* family, we examined the conservation of M^pro^ active sites from α-CoVs (*n=*4), β-CoVs (*n=*7, including SARS-CoV-2) and gamma-coronaviruses (γ-CoVs) (*n=*1) from a structural perspective. The active site amino acid sequence (Fig. 1) and conformational differences (Fig. 2) of multiple M^pro^ enzymes were compared among the selected PDB structures (Table S1). Twenty-six amino acids were selected as active site residues because they have at least one heavy atom within 4.5Å of the common ligand PRD_002214. PRD_002214 is a Michael acceptor-based peptidomimetic inhibitor, known as N3, developed previously to target M^pro^ from multiple CoVs (21–24).Since then, this inhibitor has been used in broad CoV M^pro^ enzymatic and co-crystallographic studies, including the first reported SARS-CoV-2 M^pro^ crystallographic structure (33).

**Fig. 1.**
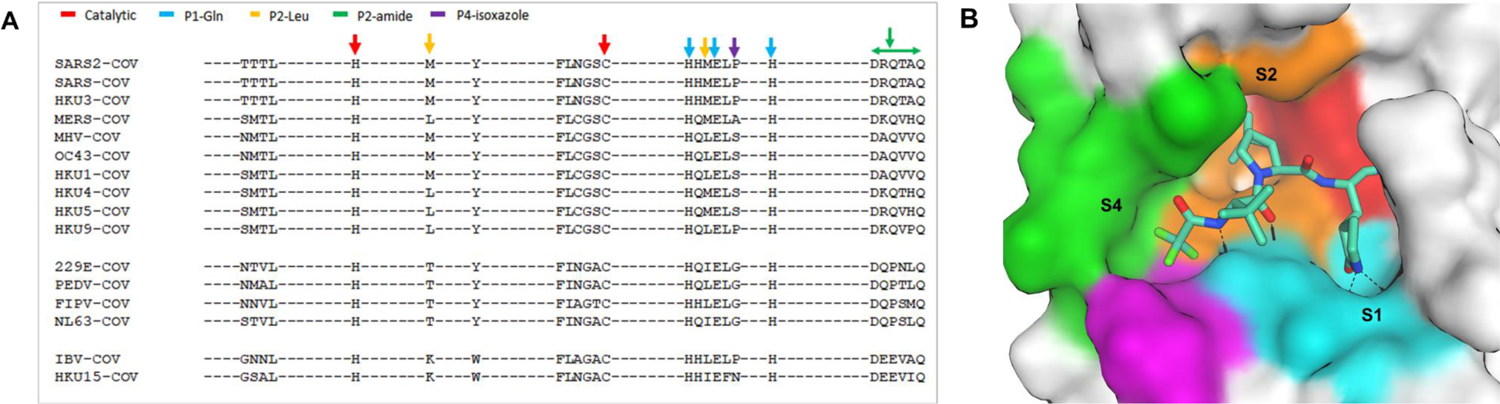
Active site conservation of CoV main proteases. A) Sequence alignment of the 26 binding site amino acids. The key amino acids are indicated by color-coded arrows based on their interaction with the inhibitor, nirmatrelvir. B) SARS-CoV-2 M^pro^ binding pocket of nirmatrelvir. The pocket surface is colored based on the inhibitor’s interaction shown in panel A.

**Fig. 2.**
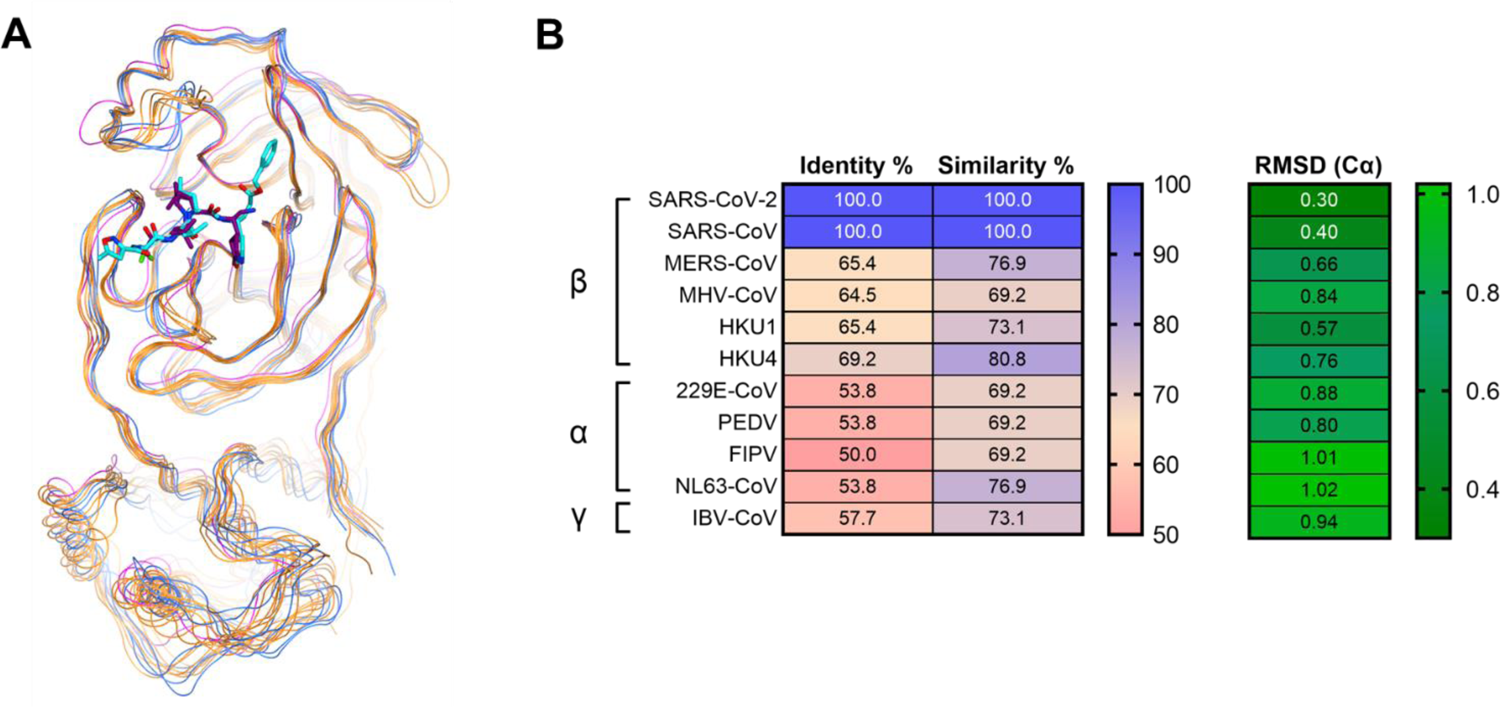
Comparison of structure and sequence identity across 12 CoV main proteases. A) Superposition of 12 CoV main proteases based on the 26 amino acid backbone heavy atoms at the active site. Proteases are represented by colored lines, β-CoV proteases in yellow lines, α-CoV proteases in blue lines and γ-CoV protease in magenta. The complete list of CoV proteases can be found in Table S1. B) Percent sequence identity, similarity and RMSD (Cα) of 26 amino acids at the nirmatrelvir binding site for β-CoVs, α-CoVs, and IBV-CoV (γ-CoV). Identity and similarity values range from 50-100 and RMSD (Cα) values range from 0.30-1.02 in their respective color mapping scales.

The sequence homology comparison of these 26 amino acid residues in M^pro^ across different CoVs is shown in Fig. 1A. The key interaction amino acids are also indicated by arrows colored by their location at the binding site (Fig. 1B). The catalytic site residues (His41 and Cys145), as well as the S1 pocket residues (His163, Glu166 and His172), which tightly interact with P1 pyrrolidinone lactamof nirmatrelvir and N3 ligands, were identical in each of the CoV M^pro^ sequences. Amino acids at S2 and S4 pockets showed slightly more diversity compared to those at S1. The S2 Met49 or Met16 residues become Leu in other β-CoV proteases or Thr in α-CoV proteases (Fig. 1A). The S4 amino acids indicated by the green arrows in Fig. 1A showed even greater diversity compared to those in S2. Though the S2 and S4 amino acids are not completely conserved across different proteases, they still share high sequence similarity. Superposition of the crystal structures of the 12 CoV M^pro^ enzymes illustrated that while they are from different genera and display varying levels of sequence identity, they were also structurally similar (Fig. 2A). This is particularly evident within the active site, where the RMSD of the structures were within 1Å (Fig. 2B). SARS-CoV-2 and SARS-CoV also shared 100% similarity and identity at the 26 active site residues (Fig. 2B). Overall, we found both the structure and sequence of the M^pro^ nirmatrelvir binding pocket were highly conserved among different CoVs.

### Mutation landscape of M^pro^ from SARS-CoV-2 genomes

An in-house annotation pipeline was developed to monitor amino acid changes in M^pro^. This pipeline enabled regular retrieval and annotation of the M^pro^ sequence of SARS-CoV-2 genomes obtained from GISAID since the beginning of the pandemic. As of January 14, 2022, 4,892,468 SARS-CoV-2 genomes collected from >250 countries were annotated and examined for mutations in the M^pro^ gene. While ∼84% of isolates share the same M^pro^ protein sequence as the reference isolate, ∼14K unique nucleotide alleles and ∼4,800 protein variants have been identified for M^pro^. The non-synonymous mutation rate (substitution/residue/year) was estimated to be 2.43E-04 for M^pro^, which is lower than RdRp (9.18E-04) and >10 fold lower than S (2.81E-03). The accumulation of amino acid changes per month were plotted for the S, RdRp, and M^pro^ proteins (Fig. 3A). Non-synonymous changes in M^pro^ remained relatively low and constant compared to RdRp and S prior to December 2021. The first rise of the non-synonymous mutation rate in S gene occurred during November through December 2020, which is consistent with emergence of the first two VOCs (Alpha and Beta). Due to the large wave of Omicron isolates collected since the end of 2021, the rate of amino acid changes in both M^pro^ and S has been increasing, with the rise for the S protein being more dramatic compared to M^pro^ and RdRp (Fig. 3B).

**Fig. 3.**
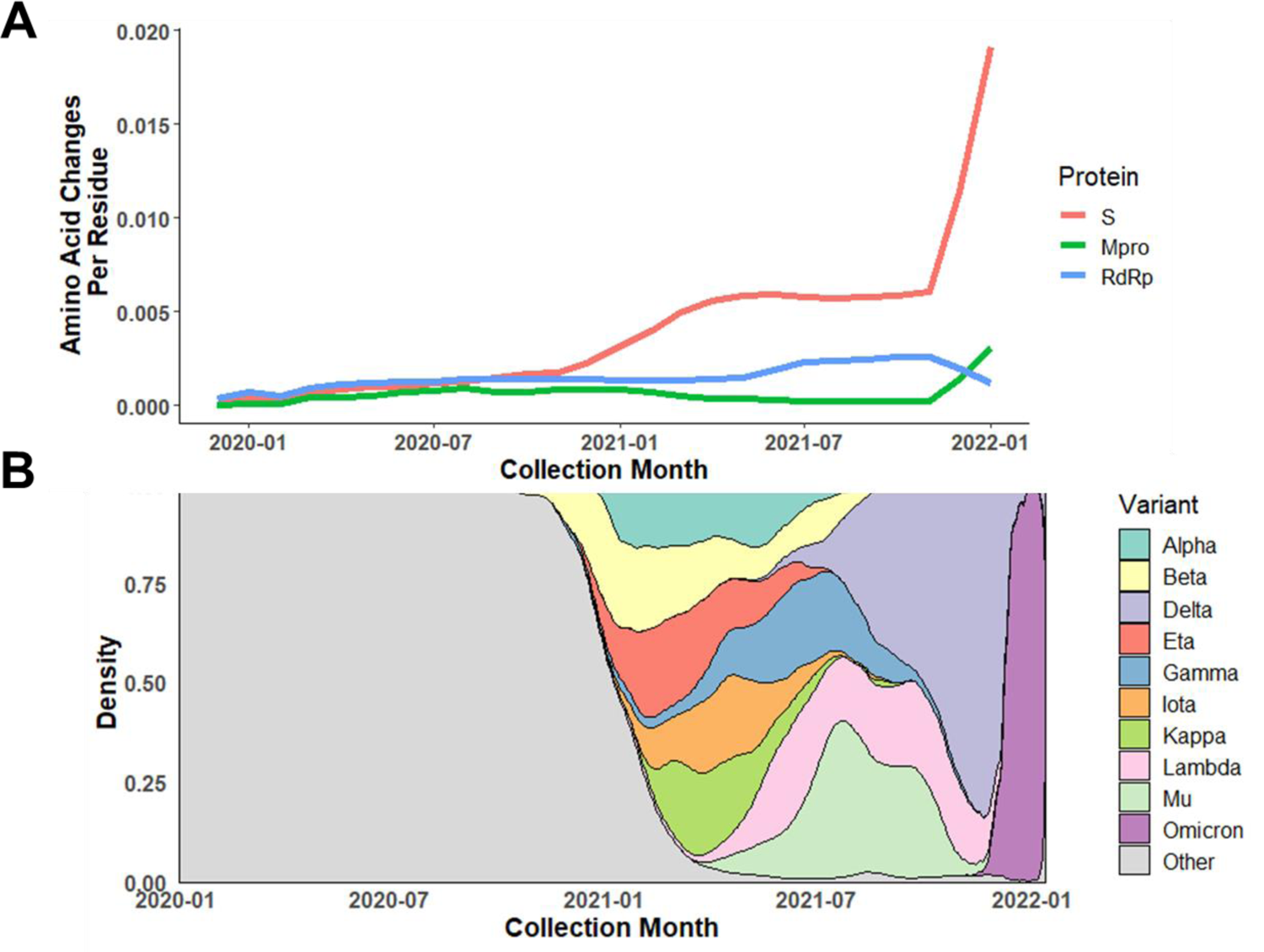
Dynamic change in amino acid mutation rate of M^pro^ compared to S protein and RdRp. A) Average amino acid changes per residue in M^pro^, S protein, and RdRp among isolates collected from January 2020 through January 2022. B) Relative distribution of VOC/VOIs based on collection date. The rapid rise in amino acid changes found in S protein and M^pro^ near the end of 2021 corresponds to the emergence and takeover of Omicron.

The key driver for the evolution of SARS-CoV-2 and numerous VOCs has primarily been adaptive amino acid changes observed in the S protein that have enabled evasion of vaccine-elicited immunity or neutralization by mAb therapeutics (34–40). Other than the selection imposed due to its essential function in viral replication, and unlike S, M^pro^ has not been subjected to vaccine or antiviral pressure to evolve. It is expected that essential function proteins like M^pro^ are under purifying (negative) selection with a signature non-synonymous-to-synonymous substitution ratio (***d*_N_/*d*_S_**) of less than 1. We conducted a selection analysis using three independent downsampled datasets of three genes, M^pro^, RdRp and S, with ∼25K sequences in each dataset. The overall mean ***d*_N_/*d*_S_** (ω) for M^pro^, RdRp and S were 0.427, 0.397, and 0.543, respectively. They were all lower than 1 and the ***d*_N_/*d*_S_** ratios for M^pro^ and RdRp were lower than that for S, suggesting that M^pro^ and RdRp were under stronger purifying selection compared to S. The nucleotide diversity (π) of M^pro^ was estimated as 6.58E-04, which was lower than that for RdRp (1.02E-3) and S (2.63E-3). Variation of the codon-based ***d*_N_/*d*_S_** ratio in M^pro^ was also examined using a Bayesian sliding window model (Fig. S1). Overall, the codon-based ***d*_N_/*d*_S_** profile was similar across three independent downsampled datasets. The mean ***d*_N_/*d*_S_** ratio across 305 codons in M^pro^ ranged from 0.117 to 0.750, except at residue 46 from one of the downsampled datasets. The regions near residues 144 and 289 had lower ***d*_N_/*d*_S_** ratios compared to other regions of the protein, indicating that amino acid changes in these regions were not favored, and implying that these domains might play critical roles in M^pro^ function.

From examination of the M^pro^ gene across >4.8 million SARS-CoV-2 genomes, the most prevalent mutations (>0.2% mutation frequency) were P132H, K90R, L89F, P108S, A260V, K88R and G15S (Fig. 4). P132H, with the highest frequency of 6.15%, is exclusively associated with the Omicron VOC (B.1.1.529 or BA.1/2). Prior to the enormous influx of Omicron cases, the frequency of P132H was as low as 0.012%. All prevalent M^pro^ mutations with occurrences >5,000 are listed in Table S2, together with their geographic and genetic lineage distribution. These are associated with different emergent VOC/VOIs. None of the prevalent mutations mapped to residues critical for nirmatrelvir activity (e.g., proximity of nirmatrelvir binding pocket as shown in Fig. 1, or dimerization interface, as shown in Fig. S2).

**Fig. 4.**
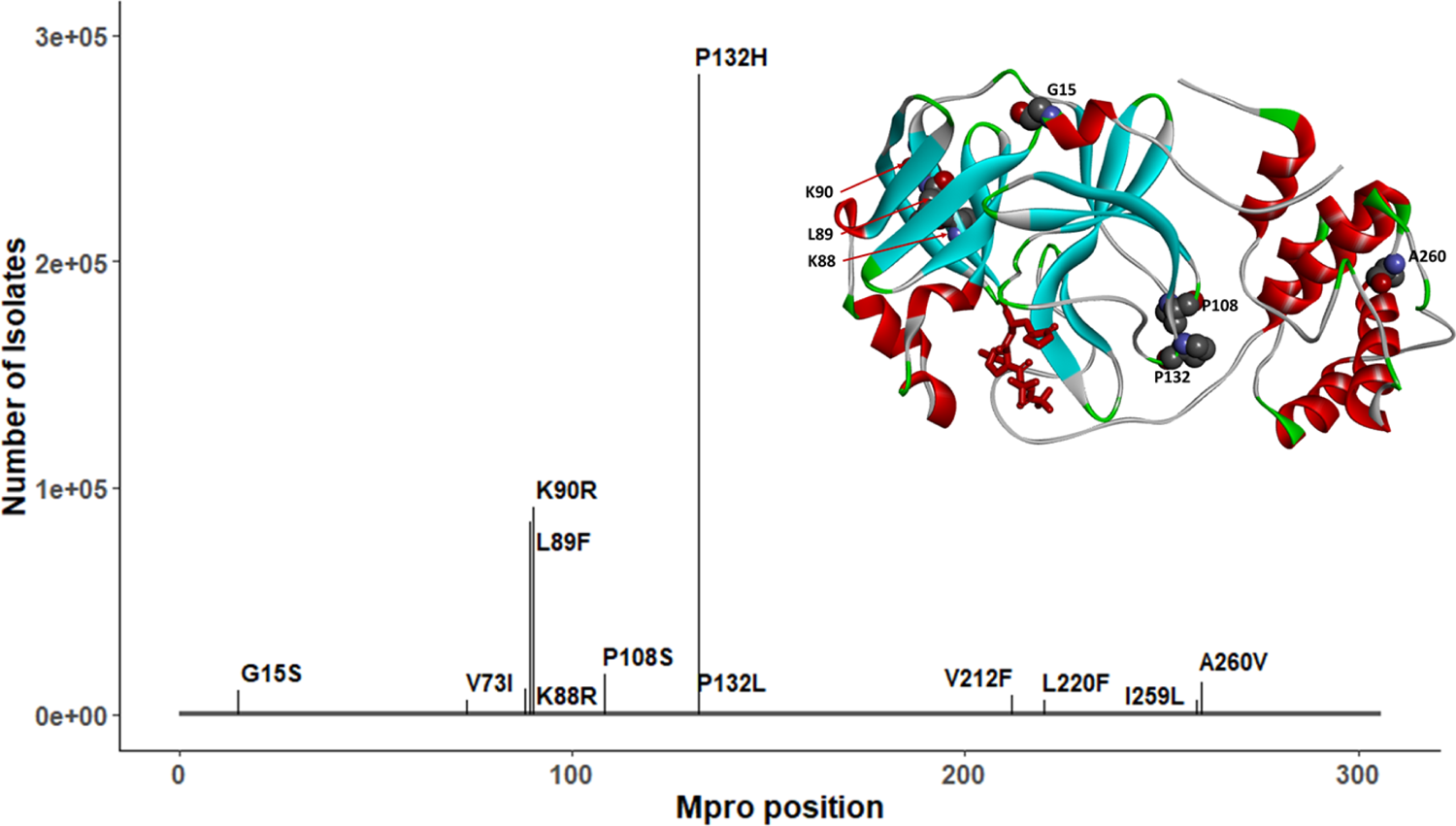
Prevalent mutations in M^pro^ and their position relative to nirmatrelvir binding. Only P132H, characteristic of the Omicron variant, exceeds 100,000 cases and no residues interact with nirmatrelvir (shown in red). The full geographic and lineage breakdown of these mutations can be found in Table S2.

### Genetic diversity of M^pro^ within variants of concern/interest (VOCs/VOIs)

In addition to the five current VOCs, two current VOIs (Lambda and Mu) and three former VOIs (Eta, Iota, and Kappa) have been identified by the WHO (7). Although emerging variants of SARS-CoV-2 are defined by accumulation of mutations in S (8), it is critical to monitor mutational changes in other viral proteins, such as M^pro^. All M^pro^ protein mutations were retrieved for each individual VOC/VOI. Aside from the Beta, Lambda and Omicron variants, the majority of isolates from each of the remaining VOCs/VOIs had M^pro^ sequences that were identical to the reference sequence (Wuhan-hu-1) (Fig. 5A). The P132H mutation was detected in >98% of Omicron isolates, whereas the most prevalent mutations in Lambda and Beta isolates were G15S and K90R, respectively (Fig. 5A). K90R is a conservative substitution and is not expected to induce changes in the 3D structure of the protease, while Gly15 is referred to as a “C’ residue” of the N-terminal α-helix (41, 42), a position with heavy preference for Gly. G15S substitution may lead to a partial decrease in the structural stability of that helix (43), although it is not likely to be detrimental to the overall protein structure.

**Fig. 5.**
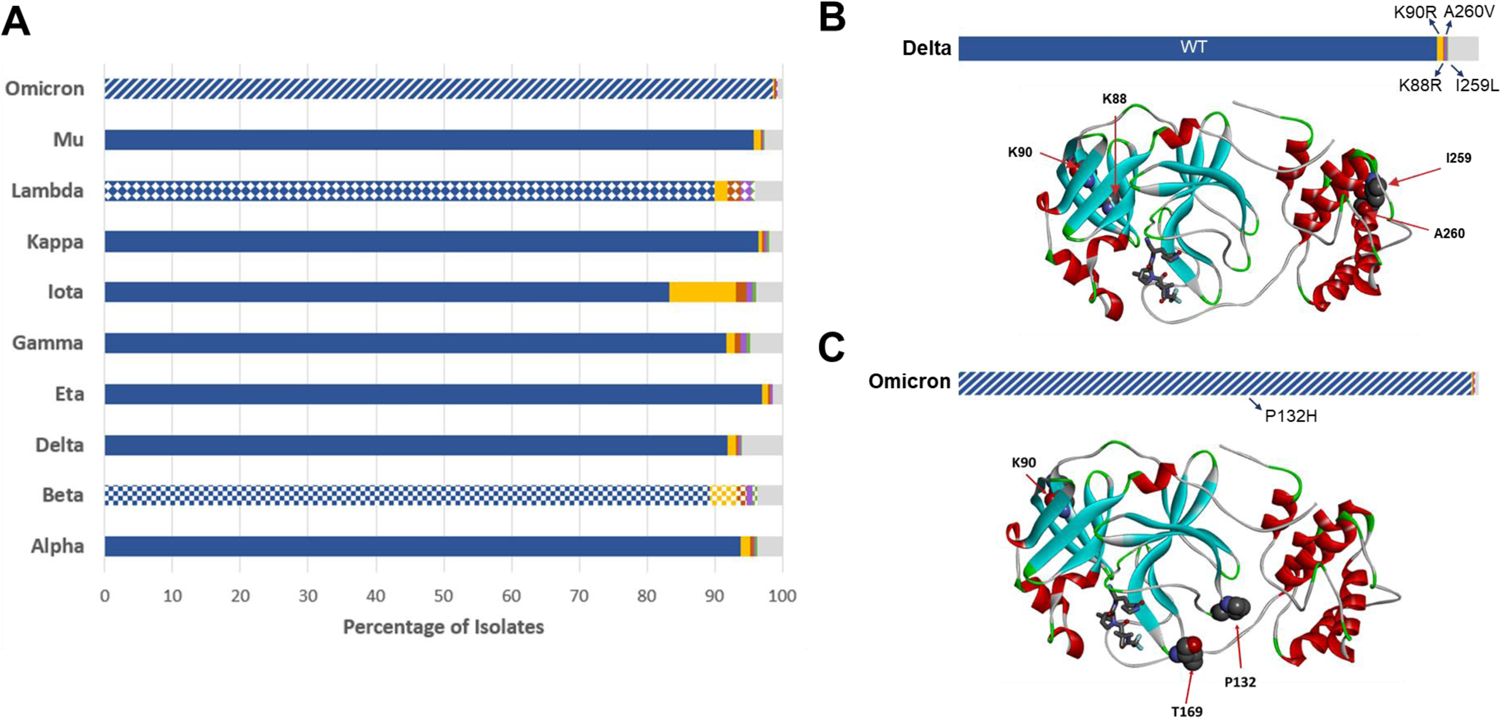
M^pro^ mutations within VOC/VOI populations. The five most prevalent sequences for each lineage are shown as colored bars (blue, gold, red, purple, green), with the cumulative remaining sequences in gray. The most prevalent sequence (blue) corresponds to the Wuhan-Hu-1 sequence (WT) and is found in all but three lineages. For these remaining lineages (Omicron, Lambda and Beta), each characteristic non-synonymous substitution is designated as a pattern: P132H (hashes), G15S (diamonds), and K90R (squares). B) Relative mutation frequency among Delta variant isolates. Position of the four most prevalent mutation sites found in this variant (K88, K90, I259, A260) are shown on the protein structure (WT). C) Relative mutation frequency among Omicron variant isolates. Position of the three most prevalent mutation sites (K90, P132, T169) are shown on the protein structure.

Prior to the Omicron surge in late 2021, Delta accounted for >90% of SARS-CoV-2 genomes submitted to GISAID (between mid-October and mid-November 2021). To investigate the potential impact of M^pro^ mutations carried by these two major VOCs on inhibitor binding interactions, we mapped the most prevalent mutation sites on the M^pro^ crystal structure with nirmatrelvir for Delta isolates (Lys88, Lys90, Ile259, Ala260; Fig. 5B) and Omicron isolates (Lys90, Pro132, Thr169; Fig. 5C). Each of these substitutions is located far from the inhibitor binding site. The most frequent M^pro^ mutation in the Omicron variant, P132H, is unlikely to affect nirmatrelvir inhibitor binding as the Pro132 residue is located within a flexible turn.

### Genetic diversity at key nirmatrelvir contact residues, cleavage sites, and the dimerization interface of M^pro^

According to the co-crystal structure of M^pro^ bound to nirmatrelvir reported earlier (11), nine key residues were identified: His41, Met49, Gly143, Cys145, His163, His164, Met165, Glu166 and Gln189 (Fig 6A). His41 and Cys145 are catalytic residues, while the remaining residues establish direct contacts with nirmatrelvir. Any changes in these residues may affect inhibitor binding. Examination of >4.8 million SARS-CoV-2 genomes illustrated that these nine residues within M^pro^ were highly conserved, with substitution frequencies of < 0.028% (Fig. 6B). Among these nine contact residues, one amino acid residue (His163) was not found to be mutated, and five residues (His41, Gly143, Cys145, His164, Glu166) were extremely conserved with ≤ six isolates identified that carry alternative amino acids. Met49, Met165, and Gln189 had more amino acid changes but still at a frequency of <0.028%.

**Fig. 6.**
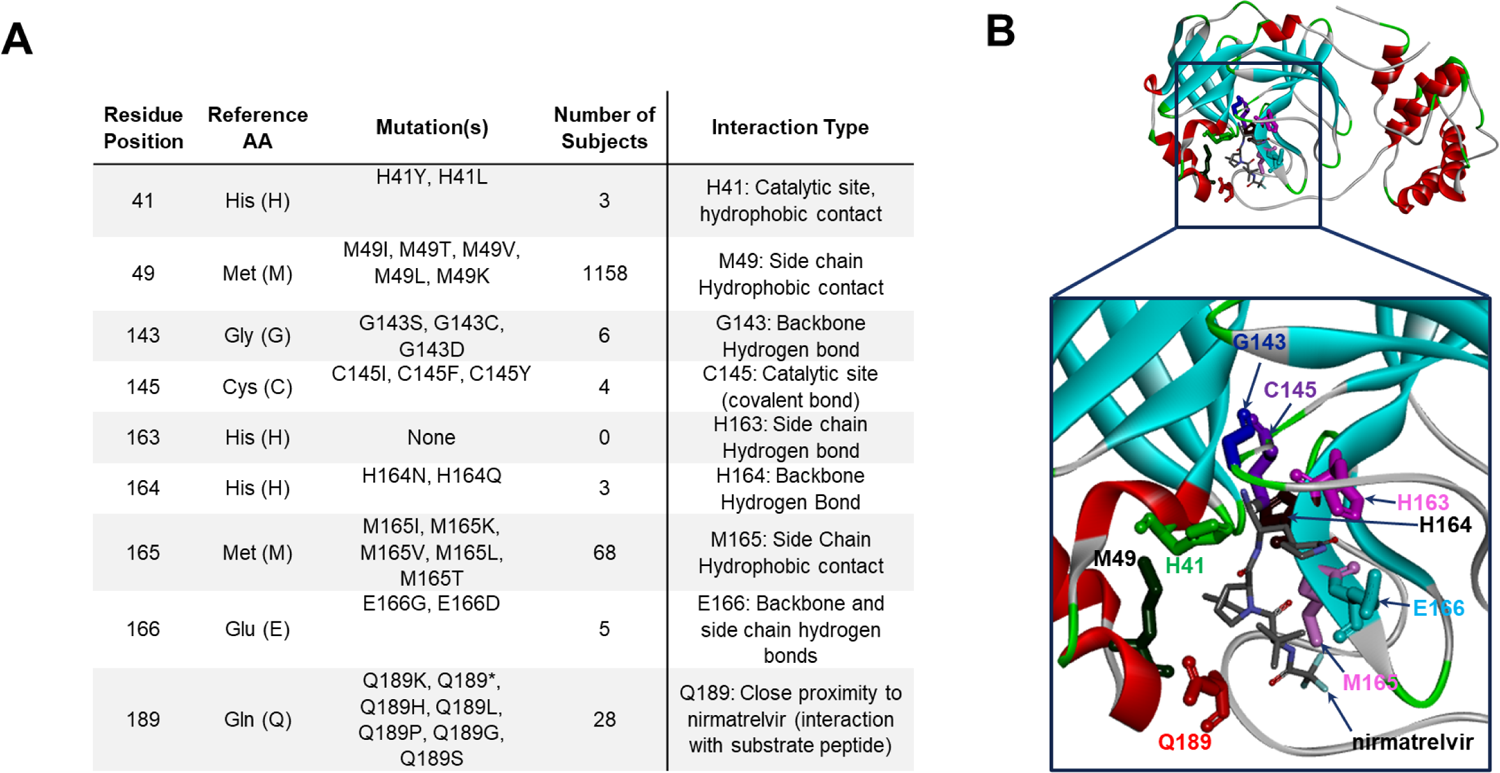
M^pro^ mutation breakdown at nirmatrelvir contact and catalytic residues. A) Mutations identified at residues directly interacting with nirmatrelvir and/or substrate peptide. B) Three-dimensional structural model of M^pro^ (PDB ID 7RFS.pdb), with residues from panel A highlighted in “stick” representation and shown in individual colors. Protein backbone is shown in ribbon representation.

Another factor that would significantly affect M^pro^ activity and catalytic efficiency is divergence from the consensus substrate recognition sequence, which always contains Gln directly upstream of the cleavage position (position P1). Preceding this (position P2) is a hydrophobic amino acid. At cleavage sites within the SARS-CoV-2 reference isolate Wuhan-hu-1, this is most commonly Leu, but some substrates contain Phe or Val at this position. The residue directly downstream of the cleavage site (P1’) is generally Ser or Ala, with Asn observed in one case. Other residues further from the cleavage position are less well conserved across target sites within SARS-CoV-2. The sequence of M^pro^ cleavage sites and neighboring residues in the reference isolate Wuhan-hu-1 (NC_045512.2) are listed in Table S3.

We investigated the mutation frequency of >4.8 million isolates at the 11 M^pro^ substrate cleavage sites and neighboring residues along ORF1ab to assess sequence conservation. In total, 445 unique amino acid changes were identified within five residues of the cleavage sites (Table S4). Despite being the most conserved amino acid among the 11 recognition sites on the Wuhan-hu-1 reference, the P1 Gln was not the most conserved residue among the examined isolates. Rather, both the P2 and P1’ positions had fewer mutations overall. In total, 7,282 instances of substitution at position P1 were observed with >98% of those cases being Gln to His (Table S4). Over 5,000 cases of this mutation were at the Nsp8-Nsp9 junction, with no more than 1,000 changes from the Gln consensus at P1 detected at any of the other 10 cleavage sites (Table S4). Consistent with the role of a hydrophobic residue at P2, ∼95% of the 4,019 amino acid changes at this position were to Leu, Ile, Val, and Phe. Meanwhile, out of 5,914 mutations at P1’, the most common was Ala to Ser – the two amino acids generally found at this position across cleavage sites. Aside from the downstream P3’ and P5’ positions, all other positions within five residues of the cleavage site had a greater incidence of mutations than positions P1, P2, and P1’ (Table S4).

M^pro^ dimerization is critical for enzyme function and the strength of the interprotomer contact can directly affect protease activity (44–47). Given the importance of dimerization, we performed analysis of amino acid residue conservation at this interface (Table 1). That interface is formed by the N-terminal tail of each protomer inserted between the two subunits of the enzyme, with many residues forming a complex network of interactions. Seventeen residues predicted to impact dimerization through interaction with one another were identified (Fig. S2). As predicted from the dimerization requirement for enzyme activity, these residues were also highly conserved with a mutation frequency of <0.11% across the >4.8 million SARS-CoV-2 genomes examined (Table 1). No substitutions were detected at Glu290 and six other residues (Glu14, Tyr126, Ser139, Glu166, Leu286 and Gln299) displayed extreme conservation with less than six instances of alternative amino acids. Residue Ala285 had the largest diversity among amino acids within the dimerization motif, although still only at a frequency of ∼0.03%.

## Discussion

For the first time, pathogen population genomics has been applied in real time to track emerging SARS-CoV-2 variants and guide the public health response to the pandemic (18). We have developed an analysis workflow to routinely annotate M^pro^ sequences and other regions of interest through genotypic surveillance. Utilizing a dataset of nearly 4.9 million SARS-CoV-2 genomes in GISAID, our analysis of the M^pro^ mutational landscape revealed that pre-existing mutations at residues interacting with nirmatrelvir, as well as at the cleavage junctions and dimerization interface, that may contribute to drug resistance were rare. The distance of the nine contact/catalytic sites to nirmatrelvir are all less than 4 Å. Notably, among the residues with key ligand interaction, only two residues (Met49 and Met165) were more frequently changed compared to others with a hydrogen bond or near the catalytic active site. Met49 and Met165 make side-chain hydrophobic contacts to the inhibitor, especially for residue Met49, which has the largest number of occurrences (*n=*1,098) among all close contact sites examined herein. It is likely that Ile at this position is acceptable since Met and Ile side chains are similar in shape and polarity, as discussed previously (48).

The considerable degree of structural similarity at the M^pro^ nirmatrelvir binding pocket across the different groups of CoVs may explain the consistent broad biochemical potency of nirmatrelvir against multiple CoVs, including SARS-CoV, Middle Eastern respiratory syndrome (MERS)-CoV, murine hepatitis virus (MHV), OC43, HKU1, 229E, NL63, and IBV proteases, as reported previously (11). In addition to the residues forming nirmatrelvir binding sites, variation in residues at the M^pro^ dimer interface was also monitored, as self-association is critical for protease activity. Although not all residues at the interface have been proven to be functionally important, it is conceivable that amino acid substitutions at positions that are spatially close to each other may introduce favorable or unfavorable interactions. In turn, this could result in changes in subunit association and, correspondingly, an impact on enzyme activity and/or nirmatrelvir binding.

Our selection analysis on M^pro^ demonstrated that the protein is under strong purifying selection with a non-synonymous-to-synonymous mutation ratio (***d*_N_/*d*_S_**) of less than 1. This is consistent with previous observations (49). However, mutations in M^pro^ could populate quickly due to the “founder effect,” when a new variant (VOC/VOI) emerges, becomes dominant in a population, and reduces genetic variation. For example, the ancestral Omicron variant always carried the P132H mutation in M^pro^. In late 2021, P132H became the most prevalent M^pro^ mutation with its frequency rapidly jumping from 0.012% to 6.15% after the Omicron surge, although this mutation does not necessarily offer any selective advantage on viral fitness, or alter inhibitor potency of nirmatrelvir (50). As expected, nirmatrelvir maintains antiviral activity against all five VOCs and two VOIs, including Omicron, Beta and Lambda which carry P132H, K90R and G15S mutations, respectively, in M^pro^ (51–55). This, however, may change with widespread use of nirmatrelvir, which, not unlike the antibodies against the S protein, may exert selective pressure on its target, leading to a reduction of potency. We anticipate, however, that this possibility would be mitigated by the key features in the chemical design and use of Paxlovid, such as maintaining structural similarity with the native substrate of M^pro^ (11), a short treatment window (5 days), and low dose of ritonavir (100 mg) (56).

It is important to note that, though this analysis provides data on what is currently circulating, this is not a prevalence-based analysis and is biased by geographic regions that are routinely sequencing isolates, with ∼55% of submitted viral genomes originating from the United Kingdom and US. Another caveat of using GISAID datasets is that only consensus genome sequences are available. Potential emerging resistant mutations usually have low frequency (minor allele) within viral quasi-species and will not be uncovered from assembled genomic contigs. The presence of artifacts in assembled sequencing data is also expected due to inevitable errors in the sequencing process. While GISAID has implemented internal checks to flag potential errors in submitted assemblies, this does not eliminate the potential risk of misinterpreting artifacts as mutations. Nonetheless, the vast number of sequences available for analysis (>7 million SARS-CoV-2 genomes as of January 14, 2022) proved valuable in providing a comprehensive picture of the mutational landscape of M^pro^.

At present, SARS-CoV-2 continues to represent a global health threat as new variants emerge. It is essential to continue tracking M^pro^ mutations in global viral isolates, especially since nirmatrelvir, the active protease inhibitor in Paxlovid, is expected to become a widely accessible COVID-19 treatment option. However, at present, nirmatrelvir has yet to be deployed on a mass scale. Following FDA approval of remdesivir, its widespread usage in hospitals for the first year and a half of the COVID-19 pandemic has permitted analyses of known resistance mutations in viral isolates under remdesivir selection (57). Therefore, as more sampled viral isolates undergo nirmatrelvir selection, and as more sequences become available in GISAID, our analysis workflow is prepared to detect the emergence of potential escape mutations. Moving forward, genomic surveillance of M^pro^ will be needed to continuously assess risk for antiviral resistance, specifically in the context of Paxlovid treatment of patients with active SARS-CoV-2 infection.

In conclusion, results of our extensive sequence analysis across nearly 4.9 million global SARS-CoV-2 isolates, including the recently emerged Omicron variant, highlight the high genetic conservation of the M^pro^ protein. We have built a robust workflow to monitor mutational changes in nirmatrelvir contact residues, polymorphism of cleavage and dimerization sites, and M^pro^ structural differences between SARS-CoV-2 and other CoVs. As new antiviral monotherapies against SARS-CoV-2 are introduced in the coming months, the potential for drug resistance is a serious concern. The genetic stability and structural conservation of M^pro^ observed over time in SARS-CoV-2 variants suggests a minimal global risk of pre-existing resistance to nirmatrelvir. An established system to surveil real-world genomic data for emerging resistant mutations is critical as the SARS-CoV-2 virus continues to evolve under the various selective pressures imposed by humans.

## Acknowledgements

We gratefully acknowledge the authors, originating and submitting laboratories of the SARS-CoV-2 genetic sequences and metadata made available through GISAID on which this research is based. The authors also thank John D. Sims, Pfizer Inc. for his support of the BIGSdb genome database to host GISAID SARS-CoV-2 genome sequences, Charlotte Allerton and Xinjun Hou, Pfizer Inc. for critical reading of the manuscript, and Christina D’Arco, Pfizer Inc. for scientific writing assistance.

All authors met ICMJE criteria for authorship and participated in the study design and conceptualization (LH, JL, AG, QY, SP, ASA, PAL), data analysis and interpretation (LH, JL, AG, QY, SP), writing (original draft) (LH, JL, AG, QY, SP), and manuscript preparation (LH, JL, AG, QY, SP, ASA, PAL, RC, YZ). All authors approved the final version for journal submission and agree to be accountable for all aspects of the work.

This research received no specific grant from any funding agency in the public, commercial, or not-for-profit sectors. This study was solely sponsored by Pfizer, Inc. All authors disclose that they are employees of Pfizer and some of the authors are shareholders in Pfizer, Inc.

## Supplementary Material

**Fig. S1.**
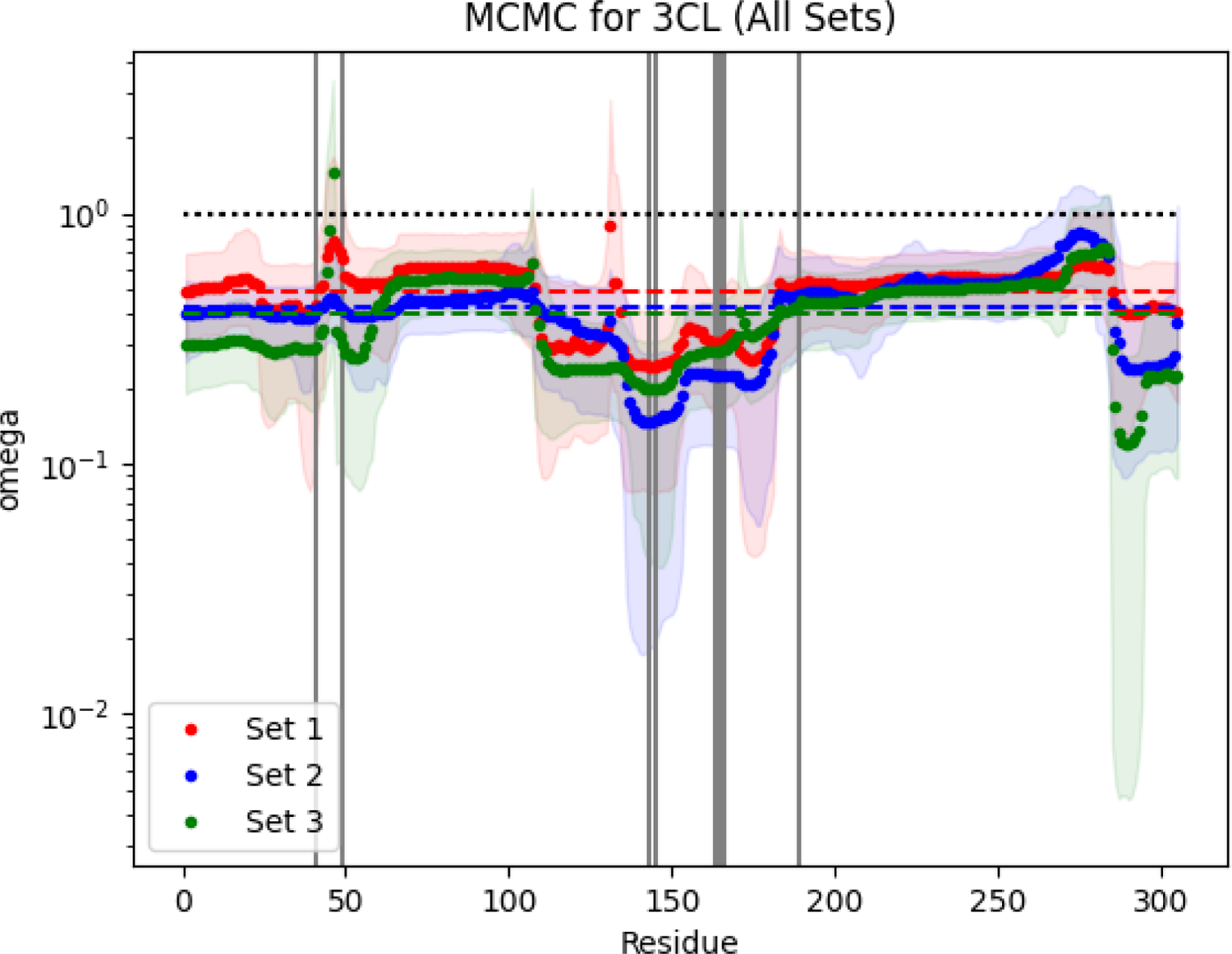
Median ***d*_N_/*d*_S_** ratio (*ω*) and 95% credibility interval along the M^pro^ gene. The ratio of non-synonymous-to-synonymous substitutions (*d*_N_/*d*_S_) was calculated using MCMC from three independent subsampling sets of the GISAID sequences (red, blue, and green) to assess the M^pro^ sequence stability. Points represent the *d*_N_/*d*_S_ at each codon and dotted lines represent the average *d*_N_/*d*_S_ for the gene. Vertical gray lines indicate codons for contact residues. Codons with *d*_N_/*d*_S_ above 1 (dotted line) indicate a greater probability for non-synonymous mutations, while those below 1 are more conserved and less favored for amino acid changes. The CI alludes to higher *d*_N_/*d*_S_ values around residues 46 and 132 (the second peak aligns with P132H).

**Fig. S2.**
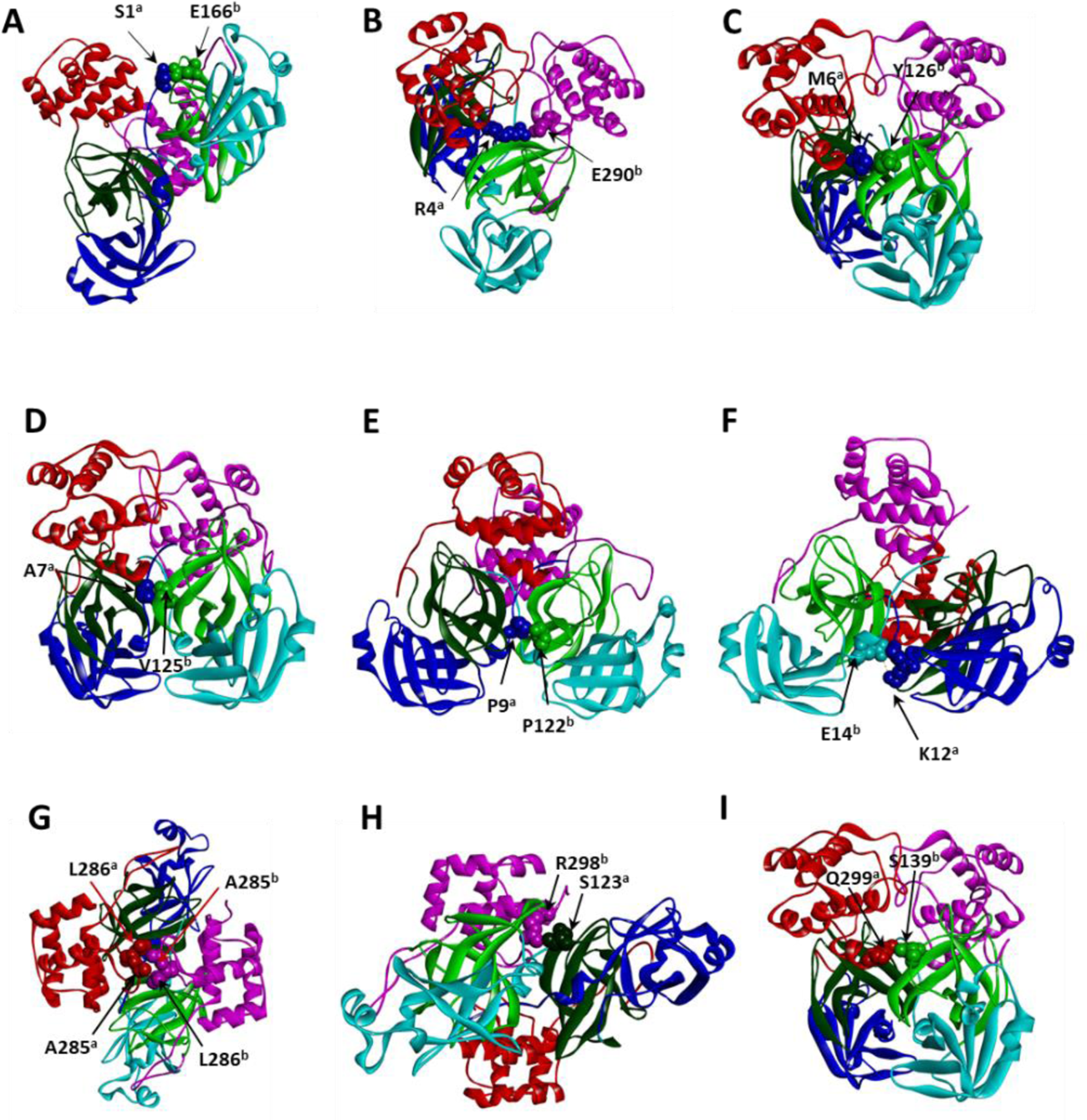
Position of dimer interface residues in M^pro^. Interface residues were identified as described in Materials and Methods. Each panel (A-I) shows only 1-2 protomer contacts for clarity. Residues involved in contact formation are shown in space-filling representation. Protein back-bone is shown as solid ribbons. Dark blue – domain I of subunit “A” (residues 1-99), light blue – domain I of subunit “B”; dark green -- domain II of subunit ‘A” (residues 100-182), light green – domain II of subunit “B”; dark red – domain III (residues 183-301) of subunit “A”, pink – domain III of subunit “B”. Figures were rendered with structural coordinates from the PDB entry 7RFS.pdb.

---

**Table S1.** List of CoV protease structures in the homology analysis.

| **Protein** | **Group** | **Pdb ID** | **Ligand** |
| --- | --- | --- | --- |
| SARS-CoV-2 main protease | beta | 7RFW | PF-07311332 |
| SARS-CoV-2 main protease | beta | 6LU7 | PRD_002214 |
| SARS-CoV-1 main protease | beta | 2AMQ | PRD_002214 |
| MERS-CoV | beta | 4RSP | PRD_002174 |
| MHV (Murine hepatitis virus) | beta | 6JIJ | PRD_002214 |
| HKU1 | beta | 3D23 | PRD_002214 |
| HKU4 (Tylonycteris bat coronavirus) | beta | 4YOG | 4F5 |
| 229E | alpha | 2ZU2 | apo |
| PEDV (Porcine epidemic diarrhea virus) | alpha | 5GWZ | PRD_002214 |
| FIPV (Feline infectious peritonitis virus) | alpha | 5EU8 | PRD_002214 |
| NL63 | alpha | 5GWY | PRD_002214 |
| IBV (infectious bronchitis virus) | gamma | 2Q6F | PRD_002214 |

**Table S2.** Geographic and lineage distribution of the most prevalent M^pro^ mutations (*n* > 5000).

| **Mutations** | **Population Frequency (%)** | **Number of Countries** | **Number of Lineages** | **Top Countries** | **Top Lineages** |
| --- | --- | --- | --- | --- | --- |
| P132H | 6.15 | 97 | 54 | UK (44.6%), US (32%), Denmark (6.7%) | BA.1 (98.1%) |
| K90R | 1.987 | 165 | 549 | US (26.3%), UK (10.2%), Denmark (9.8%) | B.1.351 (23.1%), B.1.617.2 (15%), B.1.1.7 (12.1%), AY.4 (10.4%) |
| L89F | 1.857 | 90 | 149 | US (92.7%) | B.1.2 (91.3%) |
| P108S | 0.381 | 84 | 247 | Japan (54.9%), US (15%), Switzerland (7.2%), UK (5.7%) | B.1.1.284 (52.9%), B.1.1.7 (15.1%), B.1.617.2 (7.9%) |
| A260V | 0.312 | 90 | 249 | US (29.9%), UK (22.3%), Germany (8.4%), Japan (5.8%) | B.1.617.2 (14.9%), AY.4 (14%), B.1.1.7 (12.3%), AY.44 (11.9%), AY.122 (10.4%), B.1.177 (5.2%) |
| K88R | 0.238 | 62 | 149 | UK (78.3%), US (7%) | AY.4 (75.7%), B.1.617.2 (5.3%) |
| G15S | 0.23 | 108 | 124 | UK (26.2%), Peru (10.5%), US (8.1%) | B.1.1.1 (43.3%), C.37 (10%), C.36 (8.4%), C.36.3 (5.7%) |
| V212F | 0.174 | 39 | 62 | Denmark (86.3%) | AY.4 (74.8%), AY.4.6 (12.1%), B.1.367 (8.6%) |
| I259L | 0.137 | 17 | 46 | US (84.7%), UK (13.9%) | AY.3 (95.8%) |
| L220F | 0.134 | 49 | 104 | US (49.2%), Denmark (37.5%) | AY.25 (31.1%), B.1.617.2 (24.1%), AY.122 (23%) |
| P132L | 0.133 | 72 | 174 | US (19.7%), UK (16.8%), Germany (14.2%), Netherlands (12.6%), Denmark (12.2%) | B.1.617.2 (18.4%), AY.9 (12.8%), AY.66 (8.1%), AY.9.2 (8%), AY.4 (7.8%), B.1.1.7 (5%) |
| V73I | 0.132 | 31 | 55 | US (98%) | B.1.617.2 (58.6%), AY.118 (24.2%), AY.4 (8.4%) |
| V157L | 0.128 | 59 | 126 | Denmark (52.1%), US (11.5%), UK (9.6%), Canada (8.9%), Germany (5.7%) | AY.43 (52.6%), AY.4 (8.8%), B.1.1 (8.2%), AY.4.3 (5.4%) |
| L75F | 0.116 | 78 | 203 | US (36.9%), UK (9.6%), Switzerland (7.2%), Germany (6.8%) | B.1.1.7 (35.6%), B.1.617.2 (9.6%), AY.3 (6.9%), B.1.526 (6.2%) |
| T196M | 0.112 | 52 | 110 | US (83%) | B.1.429 (37.4%), AY.44 (22.6%), B.1.234 (12%), B.1.617.2 (8.2%) |
| A191V | 0.111 | 82 | 245 | US (38.8%), UK (30.6%) | AY.4 (13.6%), B.1.617.2 (9.9%), B.1.177.20 (6.2%), AY.103 (5.9%), B.1.1.7 (5.8%), AY.25 (5%) |
| A129V | 0.11 | 71 | 177 | UK (44.9%), US (28.2%) | AY.4 (33.4%), B.1.617.2 (19.1%), AY.25 (5.6%), B.1.1.7 (5.1%) |

**Table S3.**
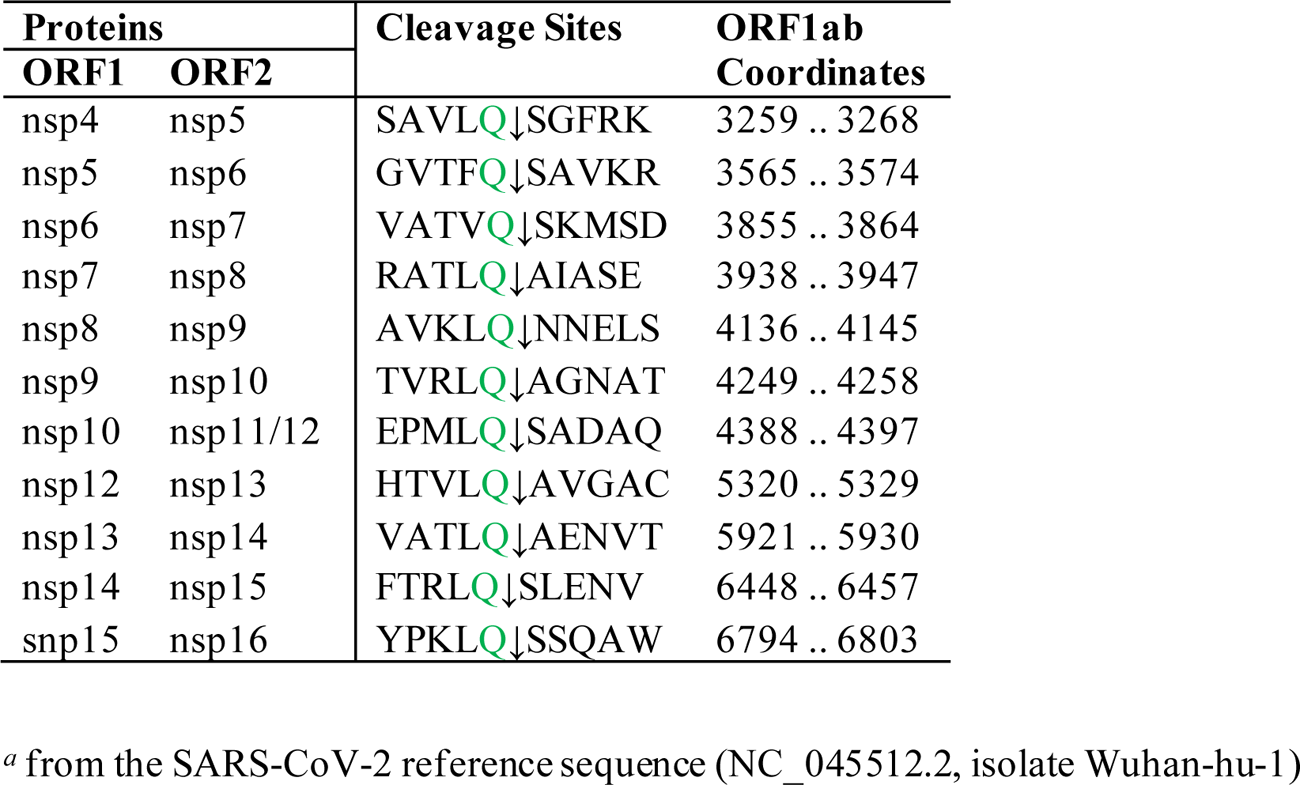
M^pro^ cleavage sites and coordinates across ORF1ab.

Table S4. Mutation frequency at M^pro^ cleavage sites and neighboring residues.

